# Experimental demonstration of tethered gene drive systems for confined population modification or suppression

**DOI:** 10.1101/2021.05.29.446308

**Authors:** Matthew Metzloff, Emily Yang, Sumit Dhole, Andrew G. Clark, Philipp W. Messer, Jackson Champer

## Abstract

Homing gene drives hold great promise for the genetic control of natural populations. However, current homing systems are capable of spreading uncontrollably between populations connected by even marginal levels of migration. This could represent a substantial sociopolitical barrier to the testing or deployment of such drives and may generally be undesirable when the objective is only local population control, such as suppression of an invasive species outside of its native range. Tethered drive systems, in which a locally confined gene drive provides the CRISPR nuclease needed for a homing drive, could provide a solution to this problem, offering the power of a homing drive and confinement of the supporting drive. Here, we demonstrate the engineering of a tethered drive system in *Drosophila*, using a regionally confined CRISPR Toxin-Antidote Recessive Embryo (TARE) drive to support modification and suppression homing drives. Each drive was able to bias inheritance in its favor, and the TARE drive was shown to spread only when released above a threshold frequency in experimental cage populations. After the TARE drive had established in the population, it facilitated the spread of a subsequently released split homing modification drive (to all individuals in the cage) and of a homing suppression drive (to its equilibrium frequency). Our results show that the tethered drive strategy is a viable and easily engineered option for providing confinement of homing drives to target populations.

## INTRODUCTION

Powerful homing gene drives can spread from low starting frequencies throughout an entire population^1–5^. However, this capability also renders such drives highly invasive since small levels of migration could facilitate their spread into any connected populations^6,7^. Though potentially desirable in some applications such as global modification or elimination of disease vectors, this could also substantially increase the social and political difficulties associated with deploying a gene drive in the field due to fears of uncontrollable spread. A gene drive that suppresses invasive species or agricultural pests would also likely raise concern if it would affect these species in their native range.

Gene drive technology has improved markedly over the past few years, and several different CRISPR-based gene drives have now been demonstrated in yeast^8–11^, flies^12,13,22–26,14–21^, mosquitoes^27–36^, and mice^37^. Most of these are homing drives, which have successfully achieved the modification and suppression of laboratory populations^21,30,36^. Several promising applications have been proposed for this technology, such as the genetic modification of *Aedes* and *Anopheles* mosquito populations by introducing genes that could reduce transmission of malaria, dengue, and other diseases^1–5^. Furthermore, gene drives could be used to directly suppress populations of disease vectors or invasive species^1–5^. Overcoming the challenge of confining a gene drive to a target population could represent an important step in bringing such approaches closer to deployment.

Several types of potentially confinable gene drives have been developed^1,38,39^. One class of especially promising candidates are CRISPR toxin-antidote (TA) gene drives. These drives will only spread in the population when introduced above a threshold frequency, which is determined by the parameters of the drive and is usually above zero if the drive has any imperfections, with some forms having nonzero introduction frequencies even in idealized form^40,41^. Below the threshold, the drive frequency will tend to decline. If migrants to connected populations cannot propel the drive above this frequency, TA drives will not spread in these populations and can thereby remain confined^40,42^. While the “migration” threshold (assuming new migrants come in each generation) will always be lower than the “introduction” threshold, the existence of one implies the existence of the other^40–42^. Recent studies have already used TA drives to successfully modify cage populations^42,43^. Despite spreading more slowly than effective homing drives and also, by design, requiring larger release sizes, such confined drives could ease concerns associated with less controllable homing drives.

The development of a confined suppression drive poses a greater challenge than confined modification drives due to the need for the drives to retain enough power to spread through the population and still provide the high genetic load required for effective population suppression. A spatially confined gene drive capable of suppression, or perhaps similarly spreading a cargo gene with a high fitness cost, would therefore be of considerable interest. Three novel systems have been suggested to this end: Toxin-Antidote Dominant Embryo (TADE) suppression drives^40,41^, daisy homing drives^44^, and tethered homing drive systems^45^. TADE suppression drives disrupt essential fertility genes and additionally target a haplolethal gene with Cas9, while the drive allele provides a rescue copy of the haplolethal gene^40,41^. These drives are capable of confined population suppression, but they require a haplolethal target gene, which can make them difficult to engineer. Many transformed individuals can be lost after embryo microinjection, since many cells would possess disrupted copies of the haplolethal gene^21^. Daisy-chain homing drives consist of a series of homing elements, with each element driving the next in the chain^44^. As the system spreads through a population, each element is lost in turn until the final element is no longer able to increase in frequency^44^. Daisy-chain drives require multiple suitable target genes to avoid resistance, and because many driving elements are required, construction is complicated^46^. In addition, with higher migration rates and lower cargo fitness costs, daisy-chain drives could be more difficult to confine^46^ and unable to keep intermediate levels of suppression for long if they fail to rapidly eliminate the population.

Tethered homing drive systems provide a potential alternative to solve some of these issues. A tethered homing drive system (Figure 1) involves the modification of a population with a confined drive, along with the spread of a homing drive that is only capable of drive activity in the presence of the confined drive^45^. Thus, a tethered system potentially combines benefits of confinement with the power of a homing drive (Table S1). Modeling suggests that such a system could be capable of confined suppression of a target population^45^. A tethered system for modification should also be able to spread cargo genes with much higher fitness costs than other confined drives.

**FIGURE 1.**
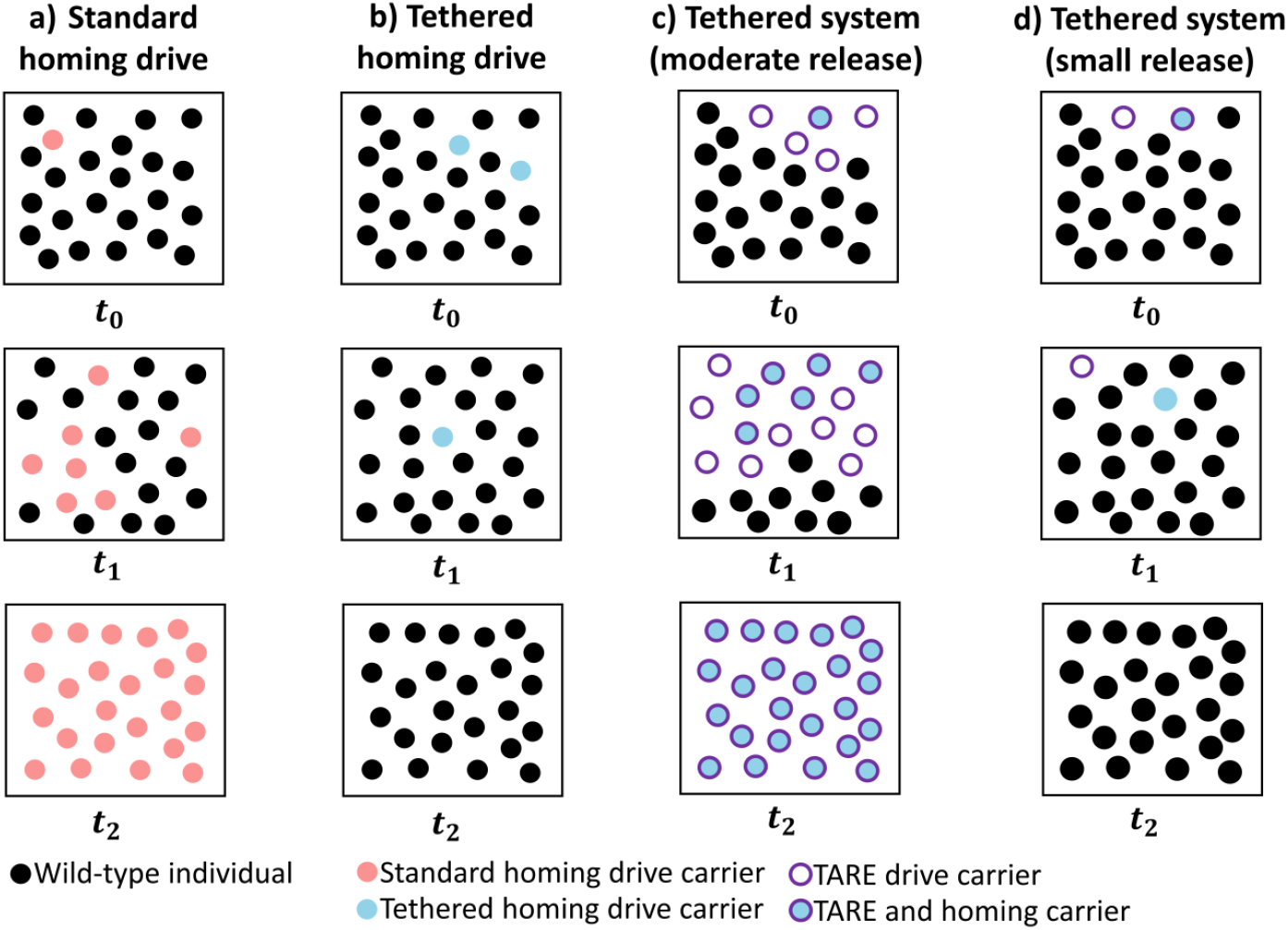
Standard and tethered homing drive comparison. Populations are shown at three different times after drive release. **a)** A standard homing drive has no threshold frequency, and it can spread to a wild-type population after a small release. **b)** A tethered homing drive cannot increase in frequency within a wild-type population since it lacks Cas9 for drive conversion. **c)** A TARE drive can spread after a moderate release, followed by a tethered homing drive after a small release. **d)** A sufficiently small introduction of TARE drive to a population will not allow the TARE drive to spread, which will in turn prevent spread of the tethered homing drive.

Here, we show that a tethered gene drive system can modify cage populations of *Drosophila melanogaster*. We use a regionally confinable Toxin-Antidote Recessive Embryo (TARE) drive targeting an essential but haplosufficient gene. This drive already contains the necessary Cas9 gene for a tethered homing drive, making it particularly suitable for use in a tethered system. We then test two split homing drives in the TARE-modified populations: The first is a modification drive targeting a haplolethal gene with two gRNAs. The second is a suppression drive targeting a haplosufficient but essential female fertility gene with four gRNAs. The TARE drive spread successfully when released well above its introduction threshold frequency and then was able to provide a Cas9 source for the homing drives, which increased in frequency according to their expected behavior.

## METHODS

### Plasmid construction

The starting plasmid EGDh2 was constructed previously^42^ (see Supplementary Information). Restriction enzymes for plasmid digestion, Q5 Hot Start DNA Polymerase for PCR, and Assembly Master Mix for Gibson assembly were acquired from New England Biolabs. Oligonucleotides were obtained from Integrated DNA Technologies. 5-α competent *Escherichia coli* from New England Biolabs and ZymoPure Midiprep kit from Zymo Research were used to transform and purify plasmids. See Supplementary Information for a list of DNA fragments, plasmids, primers, and restriction enzymes used for cloning.

### Generation of transgenic lines

Injections were conducted by Rainbow Transgenic Flies. The donor plasmid TAREhNU2G (507 ng/μL) was injected along with plasmid EGDhg2t (149 ng/μL), which provided gRNAs for transformation and was constructed previously^42^, and pBS-Hsp70-Cas9 (442 ng/μL, Addgene plasmid #45945), which provided Cas9. A 10 mM Tris-HCl, 100 μM EDTA solution at pH 8.5 was used for the injection. Both lines containing split homing modification^21^ and suppression^47^ drives were generated in previous studies.

### Genotypes and phenotypes

TARE drive carriers are indicated by expression of EGFP driven by the 3xP3 promoter, which is highly visible in the eyes of *w*^*1118*^ flies. Flies carrying either of the homing drives are similarly marked by 3xP3-DsRed, which can be easily distinguished from EGFP. For phenotyping, flies were frozen, and scored for red and green fluorescence in the eyes using the NIGHTSEA system.

### Fly rearing

Flies were reared at 25 °C with a 14/10 h day/night cycle. Bloomington Standard medium was provided as food. The cage study used 30×30×30 cm (Bugdorm, BD43030D) enclosures. Flies homozygous for the TARE drive were allowed to lay eggs in one or two food bottles for one day, and a proportionately higher number of *w*^*1118*^ individuals were separately allowed to lay eggs in six or seven food bottles for one day. All adults were removed, and the eight bottles were placed in one cage. Eleven days later, the old bottles were replaced with fresh food bottles, and the adult flies were left in the cage and allowed to lay eggs for one day before being removed and frozen for phenotyping. Adults emerging from the original bottles were considered to be generation zero. This 12-day cycle was repeated until the TARE drive had nearly reached fixation. Flies heterozygous for each homing drive and homozygous for the TARE drive were separately allowed to lay eggs in food bottles for one day at the same time as the cage flies were laying their eggs. The adults were removed before the bottles were placed into the cages with TARE drive bottles so that they were under the same conditions and would hatch at approximately the same time. The same 12-day cycle was repeated, with all flies phenotyped shortly after being frozen.

To prevent accidental releases of gene drive flies, all live drive-carrying flies were quarantined at the Sarkaria Arthropod Research Laboratory at Cornell University under Arthropod Containment Level 2 protocols in accordance with USDA APHIS standards. In addition, the homing drives contained no Cas9 and were incapable of drive conversion in wild-type flies. All safety standards were approved by the Cornell University Institutional Biosafety Committee.

### Phenotype data analysis

To calculate drive parameters, we pooled all offspring from the same type of cross together and used the combined counts to calculate rates. Because offspring had different parents and were raised in separate vials, this pooling approach could potentially distort rate and error estimates. To account for such batch effects, we performed an alternate analysis similar to the one used in previous studies^20,21,42^. This involved fitting a generalized linear mixed-effects model with a binomial distribution (fit by maximum likelihood, Adaptive Gauss-Hermite Quadrature, nAGQ = 25). In this model, offspring from a single vial were considered as a separate batch, even if they had the same parents as offspring from other vials. This approach allows for variance between batches and results in slight changes to rate and error estimates. The analysis was performed using the R statistical computing environment (3.6.1) with the packages lme4 (1.1-21, https://cran.r-project.org/web/packages/lme4/index.html) and emmeans (1.4.2, https://cran.r-project.org/web/packages/emmeans/index.html). The specific R script we used for this analysis is available on Github (https://github.com/MesserLab/Binomial-Analysis). The resulting rate estimates and errors were similar to the pooled analysis and are provided in Data Sets S1-S3.

### Genotyping

To extract DNA, flies were frozen and ground in 30 μL of 10 mM Tris-HCl pH 8, 1mM EDTA, 25 mM NaCl, and 200 μg/mL recombinant proteinase K (Thermo Scientific). The mixture was incubated at 37°C for 30 minutes, then at 95°C for 5 minutes. The DNA was used as a template for PCR using Q5 Hot Start DNA Polymerase (New England Biolabs), following the manufacturer’s protocol. The region of interest containing gRNA target sites was amplified using DNA oligo primers hCut_S_F and hCut_S_R (see supplement for primer sequences). DNA fragments were isolated by gel electrophoresis, Sanger sequenced, and analyzed with ApE software (http://biologylabs.utah.edu/jorgensen/wayned/ape).

### Estimation of drive fitness

We used a maximum likelihood approach to estimate the fitness parameters for the gene drive lines. Details of this approach and its application to cage population data were previously described^48^. The fitness parameters for both homing lines used here have been previously estimated^21,47^. For the TARE line, we estimated the viability, female fecundity, and male mating success parameters and the effective population size using a model that assumes a co-dominant, multiplicative fitness effect of the TARE allele (heterozygotes were assigned fitness equal to the square root of homozygotes). The genomic site of TARE (the *h* locus) is haplosufficient, so heterozygotes that bear one wild-type and one disrupted *h* allele were assumed to have the same fitness as wildtype homozygotes. The germline and embryonic rates of cleavage of the wildtype allele by TARE were set at experimentally estimated values. Parameter values were estimated by maximizing the likelihood across all generation transitions from the four cages combined (i.e., a single estimate per parameter is generated for all four cages). The first two generational transitions for each cage were discounted because of apparent parental effects temporarily reducing the fitness of drive individuals (resulting in fewer drive carriers in the next generation than expected), though this could also be partly explained by inbreeding between drive individuals eclosing in the same food bottle. The effective population size parameter was estimated as a fraction of the census population size in each generation transition, using the average of both generations that were part of the transition. The code is available on GitHub (github.com/MesserLab/TetheredDrives).

## RESULTS

### Drive construct design

The tethered drive concept involves the release of a confined drive and an incomplete homing drive, lacking a component provided by the confined system and thus essentially confining them to the same target area. Our tethered gene drive system involves a TARE drive carrying Cas9 together with two homing drives, one to demonstrate a population modification system and one to demonstrate a population suppression system. The homing drives, both constructed and tested in previous studies^21,47^, lack Cas9 and are incapable of drive conversion in wild-type individuals. Since the tethered system is modular, any engineered split homing drives will be compatible with the TARE drive, and any other confined system containing Cas9 can provide support for the homing drives.

The TARE drive is located in the first exon of the *hairy* locus (*h*) on chromosome 3L (Figure 2A-B) and is similar to the TARE drive used to modify a cage population in a previous study^42^. It includes a recoded *h* sequence, Cas9 expressed by the *nanos* promoter, EGFP as a marker expressed by a 3xP3 promoter and SV40 3’ UTR, and two gRNAs expressed by the U6:3 promoter. Downstream from the TARE drive, the third exon of *h* is targeted and disrupted by the drive’s gRNA. This prevents copying of the whole drive by homology-directed repair. The drive works by disrupting wild-type alleles, creating recessive lethal alleles that are then removed from the population (Figure S1A, Figure S2)

**FIGURE 2.**
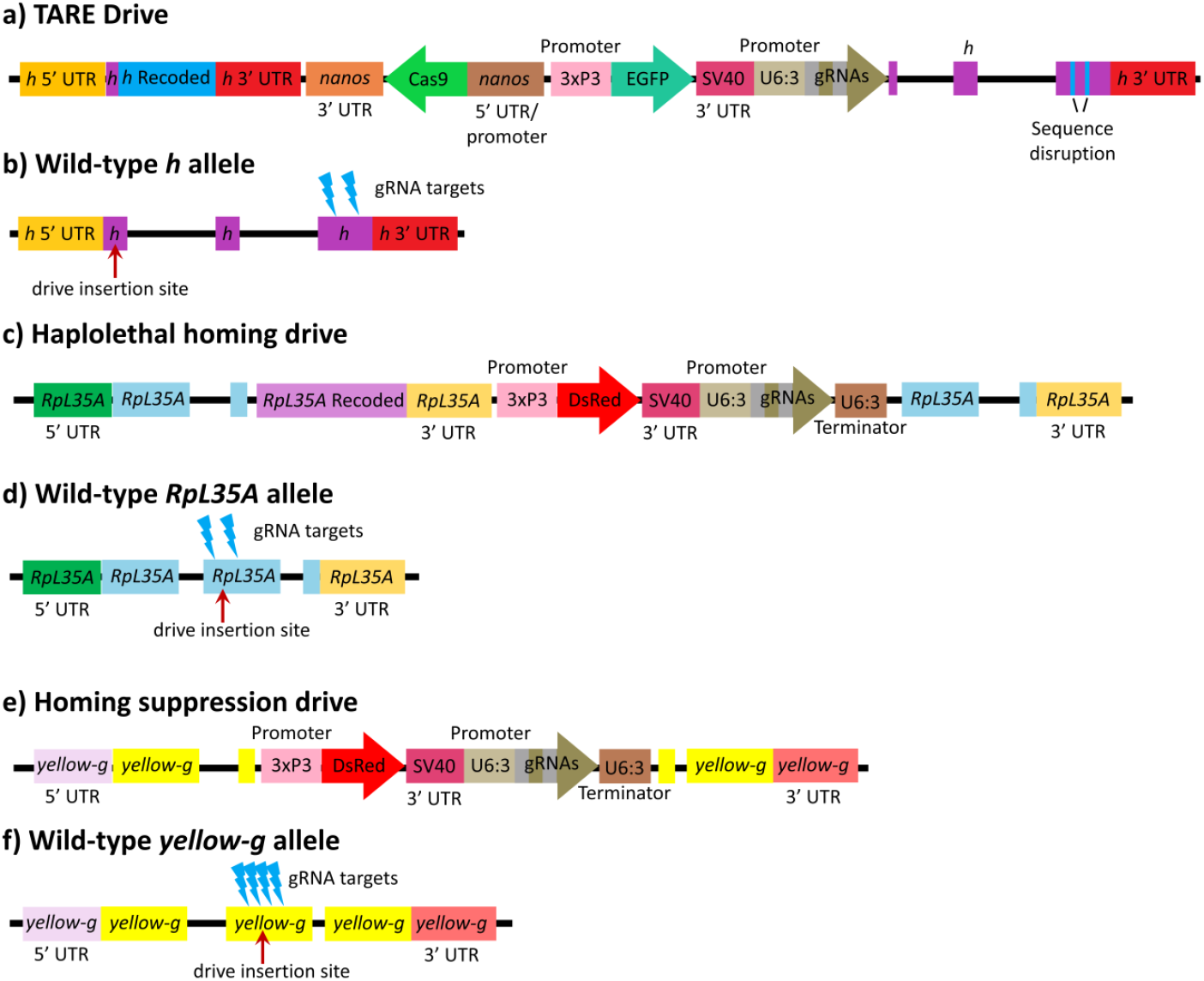
Tethered Drive Constructs. **a)** The TARE drive includes a recoded *h* sequence to provide rescue for h, Cas9 with the nanos promoter/5’UTR and 3’UTR/terminator, EGFP driven by the 3xP3 promoter with a SV40 UTR, and two gRNAs expressed by the U6:3 promoter with tRNAs to separate mature gRNAs. **b)** The TARE drive is inserted into the first exon of the wild-type *h* allele, located on chromosome 3L, and the drive targets the third coding exon with its gRNAs. **c)** The haplolethal homing drive similarly includes a recoded *RpL35A* sequence, DsRed, and two gRNAs. **d)** The haplolethal homing drive targets the second exon of the wild-type *RpL35A* allele, located on chromosome 3R. **e)** The homing suppression drive similarly includes DsRed and four gRNAs. **f)** The homing suppression drive targets the second exon of the wild-type *yellow-g* allele, located on chromosome 3L.

Our haplolethal homing drive was constructed previously^21^. In short, it is a population modification homing drive that targets the haplolethal gene *RpL35A* with two gRNAs and provides a recoded *RpL35A* as rescue (Figure 2C-D). Since the target gene is haplolethal, individuals with disrupted alleles will not be viable. Our homing suppression drive is described in detail in another study^47^. It targets a haplosufficient gene with four gRNAs (Figure 2E-F). Both homing drives are split drives and do not contain Cas9. They are only capable of drive activity in TARE-modified individuals, which express Cas9 in the germline (Figure S1B-C).

These homing drives have been engineered to reduce the incidence and impact of resistance alleles. Because the drives use multiplexed gRNAs, the formation of functional resistance alleles will be extremely rare^20^. Nonfunctional resistance alleles will lead to nonviable individuals in the haplolethal drive and will not outcompete drive alleles in the suppression drive, although the formation of large quantities of nonfunctional resistance alleles can slow the rate of spread of either drive.

### TARE drive evaluation

We crossed flies heterozygous for the TARE drive to *w*^*1118*^ flies to determine drive inheritance. Observation of the EGFP phenotype in the eyes of offspring from this cross was used to identify the presence of drive alleles (Figure 3, Data Set S1). Progeny of drive females showed 72% inheritance, a rate significantly different than Mendelian inheritance (*p* < 0.0001, Fisher’s exact test). Progeny of drive males showed 52% inheritance, which is not significantly different than Mendelian inheritance (*p* = 0.1511, Fisher’s exact test). These results were expected. Germline Cas9 activity occurs in both male and female drive carriers, creating disrupted *h* alleles, but this alone does not affect inheritance, accounting for the Mendelian rate in males. However, after fertilization, further Cas9 activity occurs in embryos from female drive carriers due to maternal deposition of Cas9 and gRNAs, even among individuals that did not receive a drive allele^40^. Any individuals with two disrupted alleles are nonviable, increasing the relative frequency of drive alleles in the progeny of drive-carrying females. The disrupted *h* alleles introduced by germline Cas9 activity in drive-carrying males remain useful when considering future generations, since they can eventually meet other disrupted alleles and then be removed from the population, further increasing the drive frequency (Figure S1A, S2).

**FIGURE 3.**
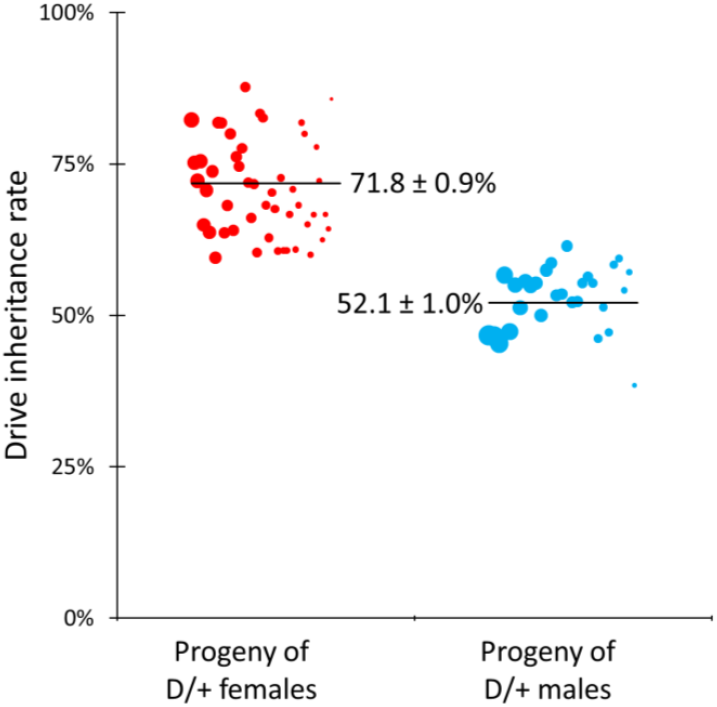
TARE Drive Inheritance. Individuals heterozygous for the TARE drive were crossed with *w*^*1118*^ individuals, and EGFP expression in progeny indicated the presence of the drive. The size of the dots is proportional to the number of adult progeny from a single drive individual. The rate estimates and standard error of the mean (SEM) are indicated. The increased inheritance in the progeny of females is due to elimination of progeny carrying no drive alleles due to maternal Cas9/gRNA deposition causing disruption in wild-type alleles inherited from the male parent. Inheritance is Mendelian in males due to lack of maternal deposition, even though *h* alleles are still disrupted in the male germline.

To determine germline and embryo cleavage rates, we crossed male and female TARE drive heterozygotes (Data Set S1). The drive inheritance rate for this cross can be used with the rate for the cross between TARE females and *w*^*1118*^ males to estimate the rates of embryo activity and germline activity. Using this approach, we calculated an embryo cleavage rate of 63.2% and a germline cleavage rate of 88.8% for the TARE drive, assuming similar germline cut rates in males and females and embryo cutting only in the progeny of females (see Data Set S1 for calculation). The germline cut rate is somewhat lower than observed in other studies of gene drive in *D. melanogaster*^13,15,16,18–21,26,42,43^ but is still sufficient to support effective population modification based on previous modeling^40^.

To further characterize and quantify the mechanism of our TARE drive, we crossed drive/wild-type heterozygotes with *w*^*1118*^ individuals. Flies were allowed to lay eggs for three days, and then transferred to new vials once per day for the next two days. In the second and third vials, eggs were counted, and pupae were counted in addition to phenotyping eclosed adults. The progeny of female drive heterozygotes had 80.7% egg-to-pupae survival, and the progeny of male drive heterozygotes had 88.2% egg-to-pupae survival (Figure 4, Data Set S1). The difference between these egg-to-pupae survival rates is significant (*p* = 0.0013, Fisher’s exact test), though slightly lower than expected (drive inheritance among progeny of females with egg counts was also slightly lower). In crosses between wild-type and drive-carrying individuals, the TARE drive uses embryo Cas9 activity to create nonviable genotypes in the progeny of females. The difference between egg-to-pupa survival rates indicates embryo Cas9 activity in the progeny of drive females, as expected.

**FIGURE 4.**
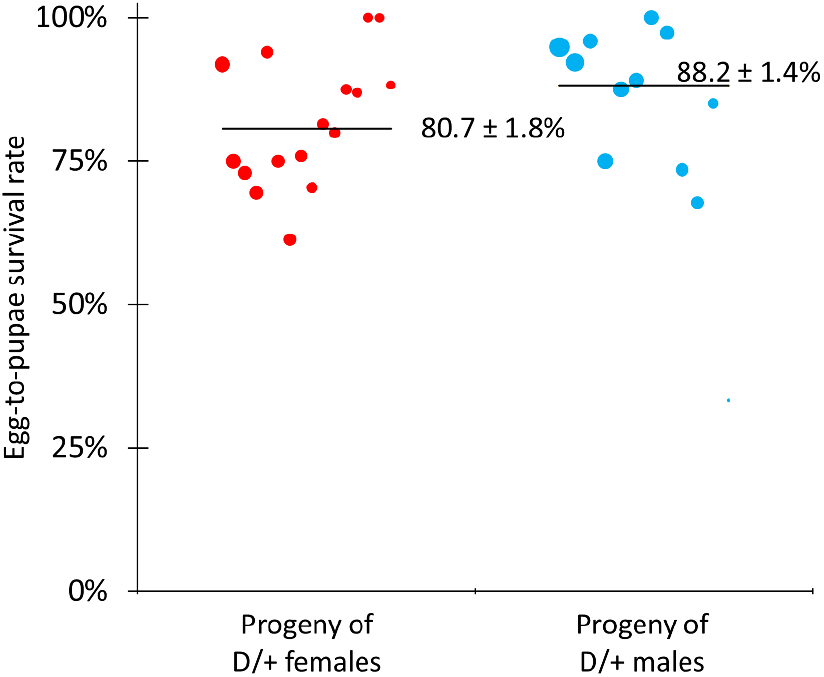
Egg-to-Pupae Viability. Individuals heterozygous for the TARE drive were crossed with *w*^*1118*^ individuals, and eggs and pupae were counted as well as eclosed adults. The size of the dots is proportional to the number of eggs from a single female. The rate estimates and standard error of the mean (SEM) are indicated. As expected, survival was reduced in egg females that inherited two disrupted *h* alleles.

### Tethered homing drive evaluation

We crossed flies doubly heterozygous for the TARE drive and one of the homing drives with *w*^*1118*^ individuals to determine drive inheritance for each homing drive. Observation of the EGFP phenotype in the eyes of offspring identified the presence of the TARE drive, and the DsRed phenotype identified either the homing suppression drive (Figure 5, Data Set S2) or the haplolethal homing drive (Figure 6, Data Set S3). In both cases, inheritance rates were lower than the rates observed in the original studies using these drive lines^21,47^ This is likely because the *nanos*-Cas9 source (part of the TARE drive) led to lower levels of Cas9 expression than the previously tested source. In addition, while the TARE drive and haplolethal homing drive are located at two distant loci, they are still on the same chromosome, and *D. melanogaster* chromosomes do not undergo crossovers in males during meiosis. This could result in suppression of the TARE drive by nonfunctional resistance allele formation from the haplolethal homing drive in this experimental cross, but the effect should be low since males with this drive have a low rate of germline resistance allele formation^21^. Egg-to-pupae survival was also determined for crosses between TARE heterozygotes and haplolethal homing drive heterozygotes (Figure S3). It was somewhat lower for progeny of TARE/homing drive females, as expected due to additional embryo nonviability resulting from the disruption of the haplolethal target gene, but it remained high for the progeny of TARE/homing drive males due to low resistance allele formation.

**FIGURE 5.**
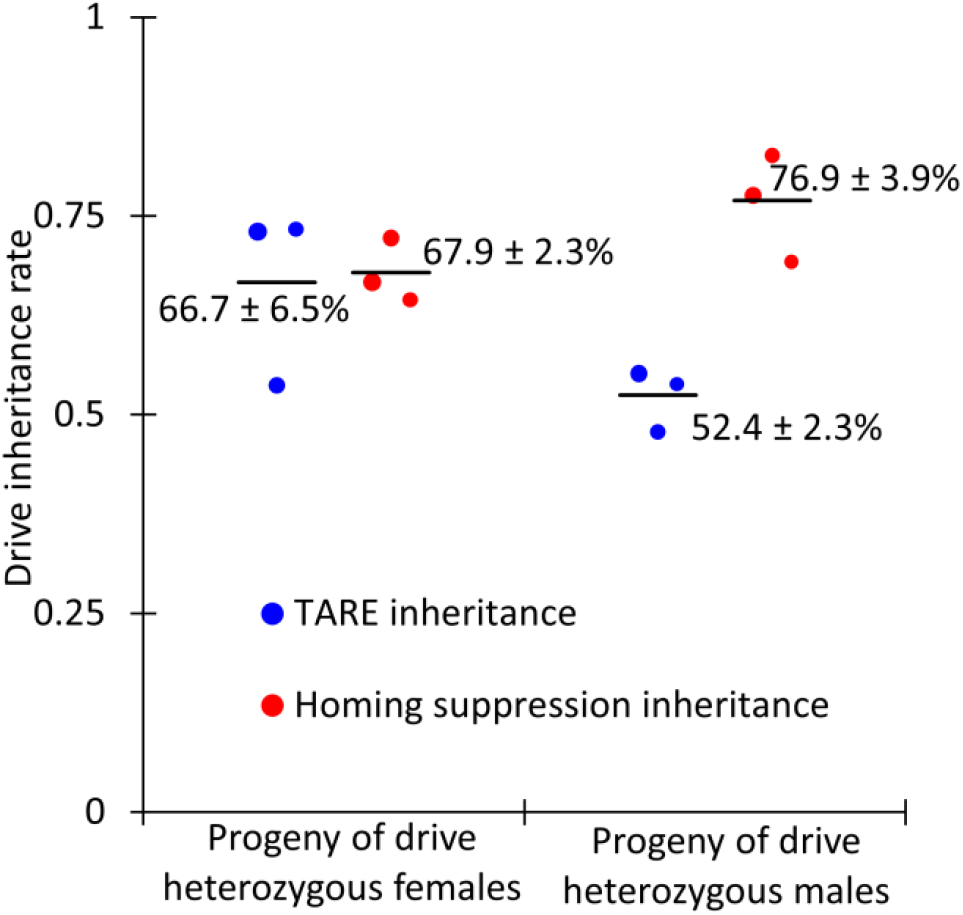
Homing Suppression Drive Inheritance. Individuals heterozygous for both the TARE drive and the homing suppression drive were crossed with *w*^*1118*^ individuals. EGFP indicated the presence of the TARE drive, and DsRed indicated the presence of the homing drive. The size of the dots is proportional to the number of adult progeny from a single drive individual. Rate estimates and SEM are indicated. The TARE drive is inherited as an increased rate only in females, but the homing drives can copy themselves, resulting in an increased inheritance rate in both sexes.

**FIGURE 6.**
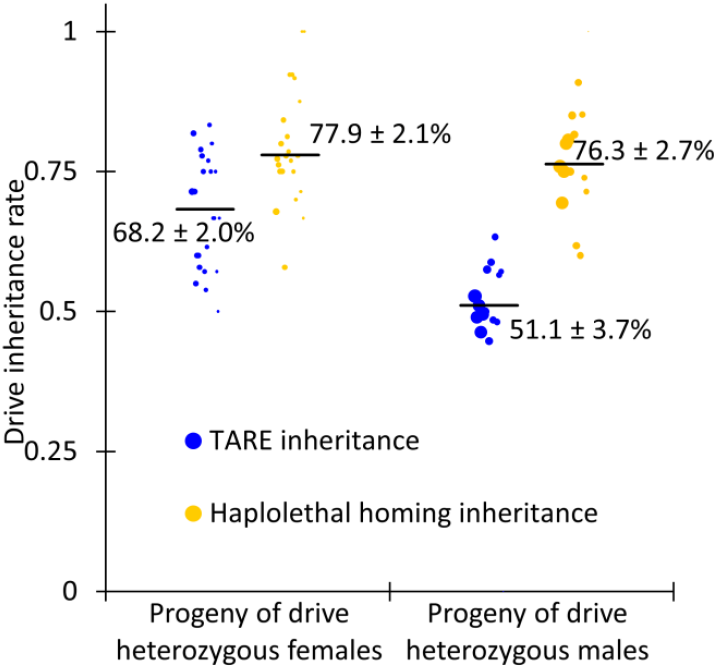
Haplolethal Homing Drive Inheritance. Individuals heterozygous for both the TARE drive and the haplolethal homing drive were crossed with *w*^*1118*^ individuals. EGFP indicated the presence of the TARE drive, and DsRed indicated the presence of the homing drive. The size of the dots is proportional to the number of adult progeny from a single drive individual. Rate estimates and SEM are indicated. The TARE drive is inherited as an increased rate only in females, but the homing drives can copy themselves, resulting in an increased inheritance rate in both sexes.

### Tethered system cage study

To assess the performance of the tethered homing drive system in large cage populations, we allowed individuals homozygous for the TARE drive to lay eggs in bottles for one day. Similarly, *w*^*1118*^ individuals were allowed to lay eggs in another set of bottles for one day. Flies were removed, and the bottles were placed in four population cages. Emerging adults were considered to be generation zero, and the TARE drive was allowed to spread in each population, with all adults phenotyped for EGFP (Figure 7, Data Set S4). In all cages, the population fluctuated, averaging 3,491 individuals in each cage (Figure S4).

**FIGURE 7.**
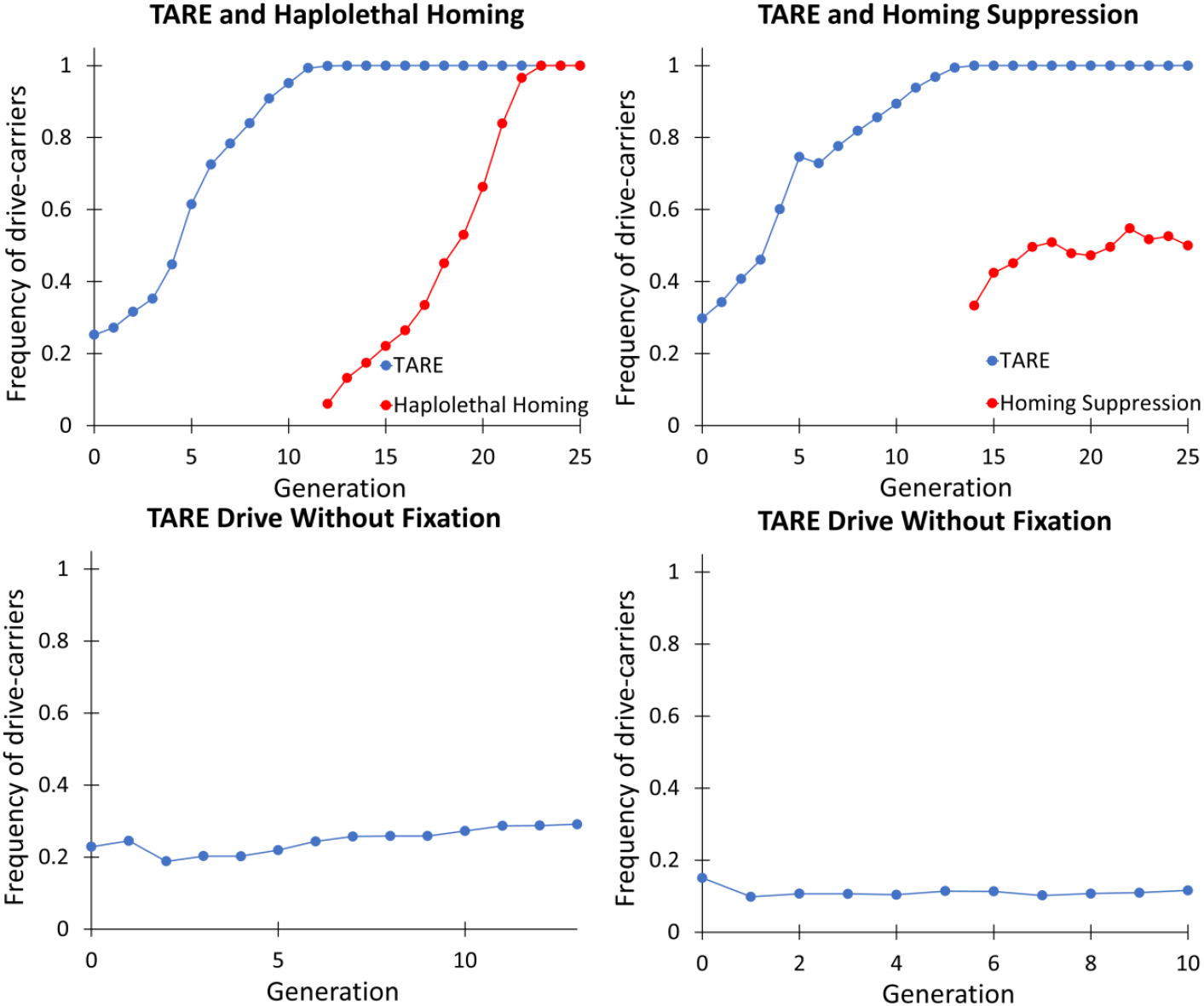
Tethered Homing System Cage Study. Individuals homozygous for the TARE drive were introduced into a population that was wild type at the drive site. The cage populations were followed for several discrete generations, each lasting twelve days, including one day of egg-laying. After the TARE drive reached fixation in two cages, individuals that were homozygous for the TARE drive and heterozygous for one of the split homing drives were released. All individuals from each generation were phenotyped for DsRed and EGFP, with positive drive carriers having either one or two drive alleles.

In one cage with a low initial release frequency, the TARE drive increased in frequency only slowly, and in another cage with a lower release frequency, the drive frequency remained constant. These two cages experienced fluctuations in frequency (possibly due to differences in the health of initially released TARE and *w*^*1118*^ individuals) before quickly reaching drive carrier frequencies of 20% and 10% (the former of which then proceeded to slowly increase, while the latter remained stable), suggesting that the introduction threshold of the TARE drive may be approximately 5-15%, a value that depends on the efficiency and fitness cost of the drive^40,42^.

With the cleavage parameters calculated based on drive inheritance (Data Set S1), the fitness cost of the drive in homozygotes was perhaps 10-15% based on previous models^40^. A maximum-likelihood based analysis^48^ indicates that drive homozygotes had approximately a 13-15% fitness cost (fitness was 0.867 compared to wild-type fitness of 1, with a 95% confidence interval of 0.785 - 0.954) if costs impacted both sexes. The fitness cost was 25-27% if only one sex was impacted (with a 95% confidence interval of 0.622 - 0.906 for female fecundity alone), and models with a single fitness parameter for female fecundity, male mating success, or offspring viability all gave similar performance (Table S2). All these models would result in an approximately 14% introduction frequency threshold with a 95% confidence interval of 3-21%. Based on the behavior of the two cages that were released close to the introduction threshold and where frequency trajectory should therefore be particularly sensitive to fitness effects, we favor the lower half of the fitness/introduction threshold range. The estimated effective population size was 3.6% of the census population size (with a 95% confidence interval of 2.4% - 5.3%), somewhat lower than similar cages analyzed by the same maximum likelihood method^21,47–49^. This is perhaps indicative of a model that does not capture all relevant factors influencing genotype frequencies^48^, and indeed, the rate of increase of the TARE drive was somewhat higher than predicted when it was at an intermediate frequency (40-70% carrier frequency). One possible explanation for this is underestimation of drive cut rates. Another is patchy distribution of eggs from drive and wild-type individuals, either between or within the eight food bottles of the cage experiment, thus creating areas with higher proportions of eggs from drive-carrying mothers. In these areas, some offspring from drive mothers would be nonviable, allowing their adjacent siblings access to more resources at the early larval stage than individuals in more crowded areas with a higher proportion of viable eggs.

In the two cages where the TARE drive reached 100% frequency, individuals homozygous for the TARE drive and heterozygous for one of the homing drives were released. The haplolethal homing drive was introduced at 6% frequency, and the homing suppression drive was introduced at 33% frequency (Figure 7, Data Set S4). The homing drives were allowed to spread for several generations, with all adults phenotyped for DsRed. The haplolethal homing drive eventually reached a frequency of 100%, while the homing suppression drive approached an apparent equilibrium frequency of somewhat over 50%. This latter result was due to low drive efficiency and a fitness cost, which prevented complete suppression and was seen in another study using the same drive^47^. The rate of increase of these homing drives was lower than in other studies that used these lines^21,47^, as expected from the somewhat lower drive conversion rate of these drives when combined with the TARE drive compared to the previously used *nanos*-Cas9 supporting element. The additional gRNAs in the TARE drive may also have somewhat reduced the cleavage activity at each site in the split homing drives due to saturation of Cas9, though this reduction was likely small^20^.

## DISCUSSION

In this study, we performed an experimental trial of the “tethered gene drive” system. This system was designed to allow a potentially confined drive system, such as a TARE drive or an underdominance system, to support a homing drive, thus allowing costly alleles to be spread with the powerful homing drive, while also preventing it from spreading to populations beyond the supporting confined drive^45^. Our demonstration with TARE followed by a tethered population modification drive^21^ was successful, though the tethered homing suppression drive did not perform as well. However, the equilibrium frequency of the suppression drive is determined by its fitness costs, drive conversion efficiency, and other characteristics of the drive itself^47^, rather than by any feature of the tethered system, which would only be expected to slightly reduce drive performance due to saturation of Cas9 with additional gRNAs in the combined system.

Since the TARE drive utilized the same rescue element, gRNA promoter, and target sites as a previous split TARE system^42^, its reduced performance as indicated by reduced drive inheritance was almost certainly due to the Cas9 element. Though both of these TARE drives had identical *nanos*-Cas9 elements (with the same orientation with respect to the 3xP3 promoter of the fluorescent element), their genomic locations were substantially different. In this study, the Cas9 gene was located within the TARE drive on chromosome 3L, rather than a separate genomic site on chromosome 2R. This led to reduced cleavage rates, particularly in the early embryo, a critical parameter that can influence drive inheritance in individual crosses. In this case, such reduced performance could potentially be useful to make the drive more confined to a target region. However, for the homing suppression element, high performance would likely be needed to achieve high enough drive conversion efficiency and genetic load for population suppression^20^, particularly in complex natural environments^50,51^. Higher expression rates of Cas9 within the TARE element could possibly be achieved by reorienting the Cas9 element within the drive allele. It would also be straightforward to choose a different site within the *h* gene, or more likely, a different target gene entirely. Essential but haplosufficient genes can likely be found in most genomes, and similar CRISPR toxin-antidote ClvR elements have already targeted other such genes with success^43,52^.

In general, the tethered system provides a highly promising strategy for confining a homing drive to a target population that is sufficiently isolated from other non-target populations. If an efficient homing drive can be made, then a suitable TARE system can likely be engineered as well. This is because a suitable homing system would have a high drive conversion rate, implying at least an equal rate of germline cleavage (which includes both the drive conversion rate and germline resistance allele formation rate for homing drives). A TARE system would be expected to still have good efficiency without embryo cutting if the germline cut rate was high (without embryo cutting, it would take only a few more generations to spread to the whole population^40^), allowing it to utilize the same promoter. While TARE would additionally need a rescue element, several studies have already shown that their engineering is feasible in flies^21,42,43,52^ and mosquitoes^36^. Furthermore, a TARE drive could be engineered to have an additional CRISPR nuclease with a different promoter and different gRNA specificity^53–55^, allowing germline-restricted CRISPR cleavage for the homing element and both germline and embryo cleavage for the TARE element. This could increase the efficiency of the homing drive, which generally suffers from embryo cutting. However, modeling indicates that a homing drive with high drive conversion efficiency or lacking significant fitness costs would still be able to tolerate high embryo cut rates, as long as functional resistance alleles did not form^20,56,57^.

For more stringent confinement than what can be provided by a standard TARE drive, other CRISPR toxin-antidote systems with higher introduction thresholds could likely be engineered with similar ease, such as 2-locus TARE systems (Table S1)^41^. This additional confinement could provide a buffer against an evolved reduction in fitness costs or high variation in migration levels that could otherwise occasionally result in a drive exceeding its introduction threshold in a non-target population. However, in situations where an efficient homing drive could not be engineered due to difficulty in achieving a high enough rate of drive conversion by homology-directed repair after germline cleavage, then TADE suppression systems^40,41^ may present a useful alternative option for confined suppression if targeting of haplolethal genes could be successfully engineered.

This study experimentally demonstrated the feasibility of using tethered homing gene drives in cage populations based on CRISPR toxin-antidote drive systems. Such systems can allow for the power of homing drives to be limited to target populations. This could be useful and even necessary when dealing with invasive species outside their native range, when sociopolitical considerations demand a limited drive, or even during an initial testing phase of a homing gene drive system.

## ACKNOWLEDGEMENTS

This study was supported by the National Institutes of Health awards R21AI130635 to JC, AGC, and PWM, award F32AI138476 to JC, and award R01GM127418 to PWM. We thank Jim Bull and two anonymous reviewers for their suggestions, which helped improve the quality of our manuscript.

## SUPPLEMENTARY INFORMATION

The following table shows the DNA fragments used for Gibson Assembly of the plasmid. PCR products are shown with the oligonucleotide primer pair used, and plasmid digest is shown with the restriction enzymes used.

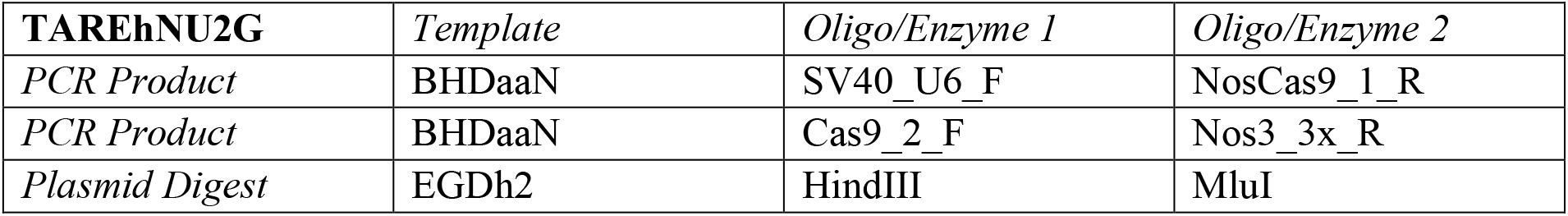

### Construction primers

Cas9_2_F: AAACAGCTCAAGAGGCGCC

Nos3_3x_R: TTCAATTAGAGCTAATTCAATTAGGATCCAAGCTTTCCTTCCTGGCCCTTTTCGA

NosCas9_1_R: TCCTGTATATCGGCGCCTCTT

SV40_U6_F: GGAGCAATCACAGGTGAGCAAAAAAACGCGTTAAGATACATTGATGAGTTTGGACAAACC

### Sequencing primers

EGFP_S_F: AGCGCACCATCTTCTTCAAGG

h3utr_S_F: AAGGACCTTCATCAGACGCAC

hCut_S_F: CCAAATTGGAAAAGGCCGACA

hCut_S_R: AACATGGGTTGCTGTTGTGC

**FIGURE S1.**
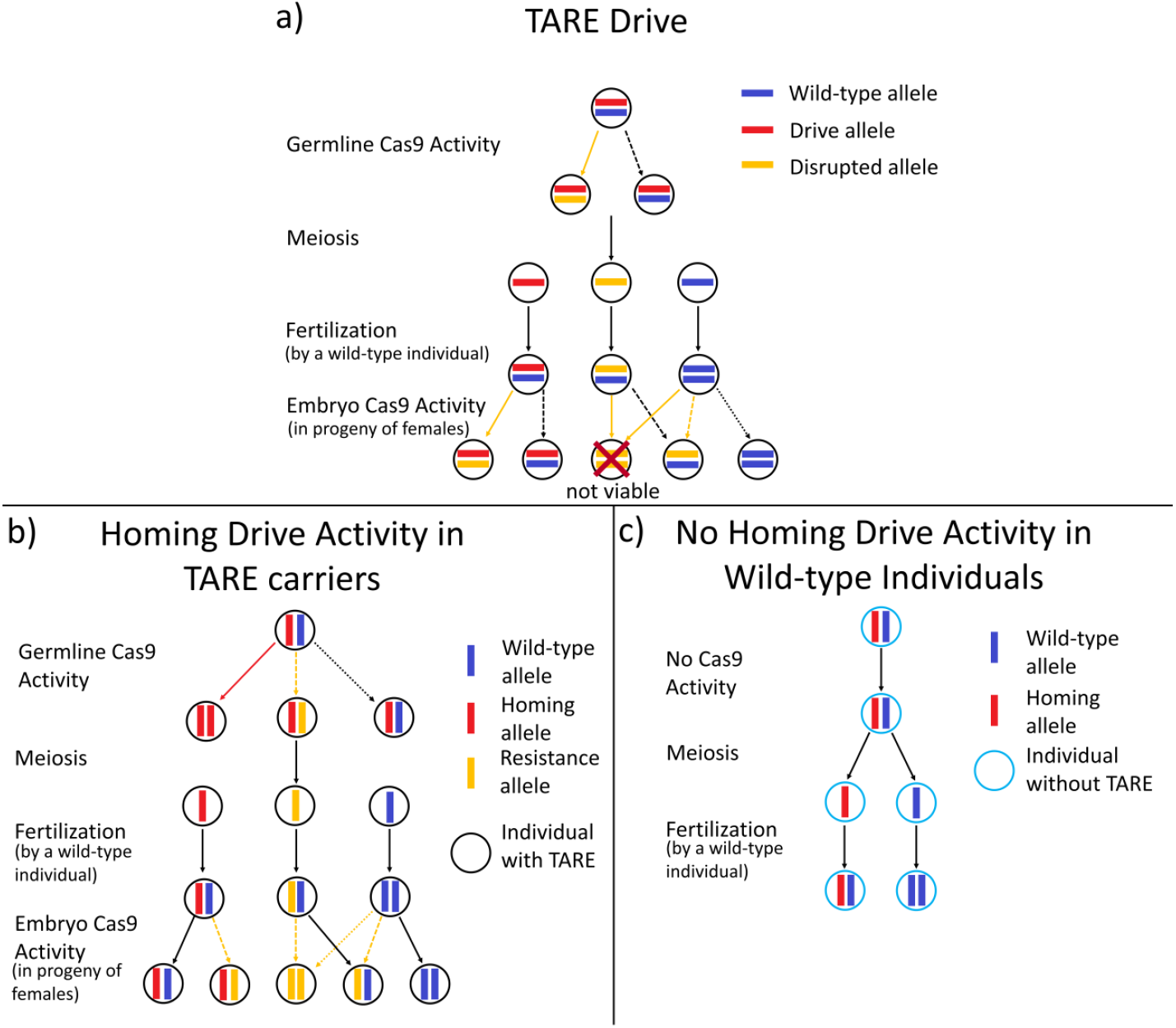
Tethered Drive Mechanism. **a)** A regionally confined TARE drive is first released. Cas9 activity occurs in the germline of drive-carrying individuals, disrupting wild-type alleles. Then, embryo Cas9 activity occurs in the progeny of females. Individuals with two disrupted alleles are not viable. Eventually, the TARE drive will spread in the target population. Common events are indicated with solid arrows, and dashed arrows indicate less common events. **b)** A homing drive is released into the population of individuals with the TARE drive (or another confined drive with Cas9). Cas9 activity leads to drive conversion by homology-directed repair or to resistance alleles (which can be removed in a manner dependent on the homing drive’s target gene). **c)** Homing drive activity cannot occur in individuals that do not express Cas9, thus confining it to the population that contains the TARE drive.

**FIGURE S2.**
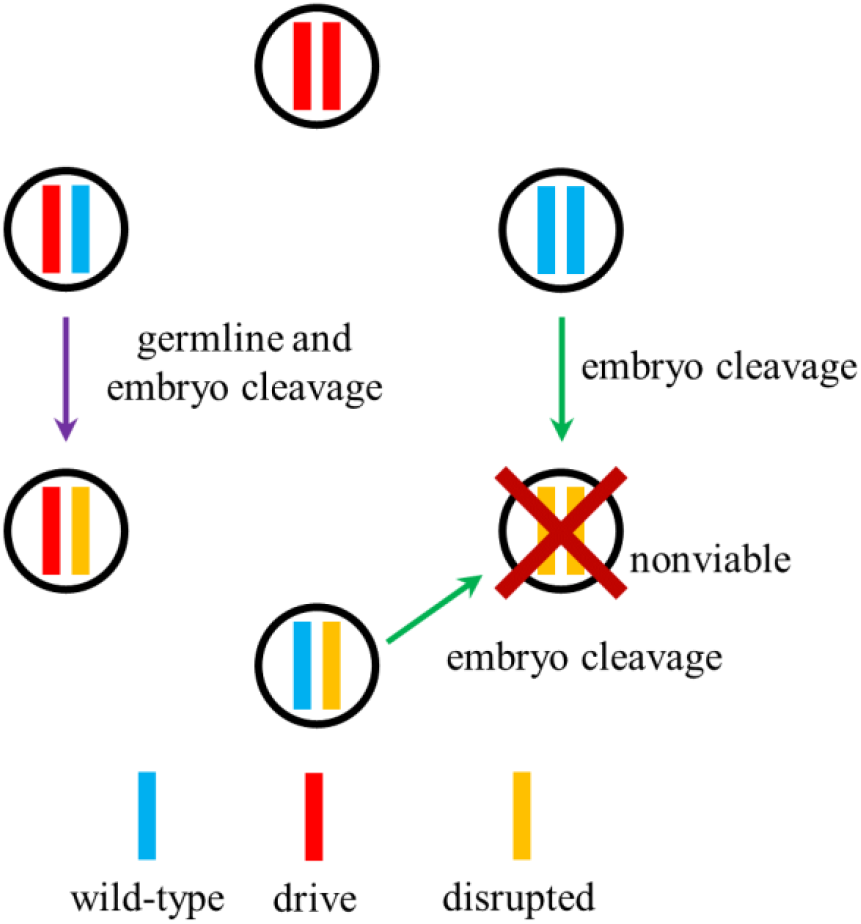
Relations between genotypes in the TARE drive. There are six possible genotypes for the TARE drive, representing combinations between drive, disrupted, and wild-type alleles. Drive/wild-type germline cells are usually converted to drive/disrupted heterozygotes in both males and females. Additionally, any wild-type allele in any genotype can be converted to a disrupted allele due to embryo Cas9 activity if the mother carried a drive allele. Individuals that are homozygous for disrupted alleles are nonviable. Thus, wild-type alleles in a population will tend to be converted to disrupted alleles, which are often nonviable, thus increasing the frequency of drive alleles in the population.

**FIGURE S3.**
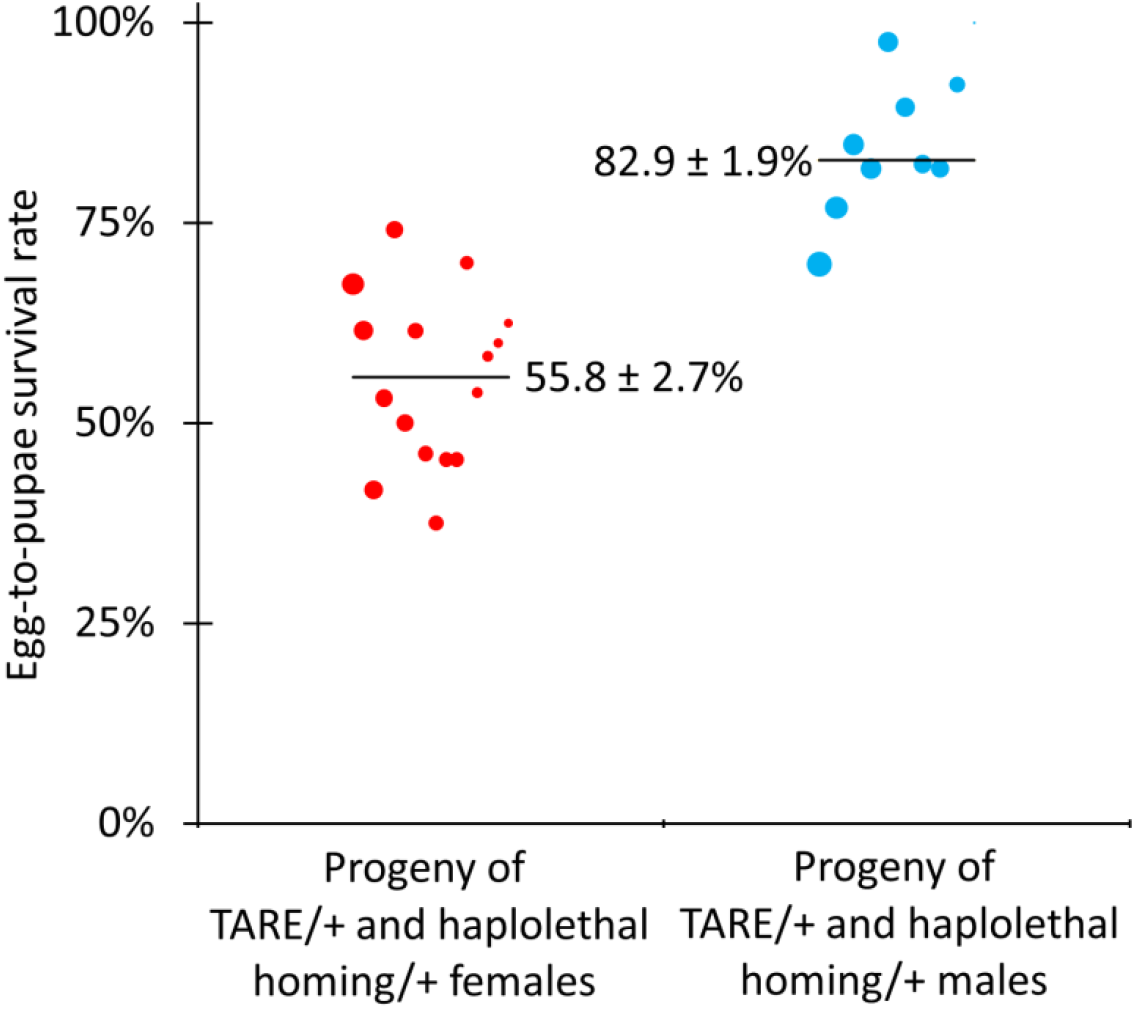
Egg-to-Pupae Viability for Haplolethal Homing Drive. Individuals heterozygous for the TARE drive and the haplolethal homing drive were crossed with *w*^*1118*^ individuals, and eggs and pupae were counted as well as eclosed adults. The size of the dots is proportional to the number of eggs from a single female. Rate estimates and SEM are indicated.

**FIGURE S4.**
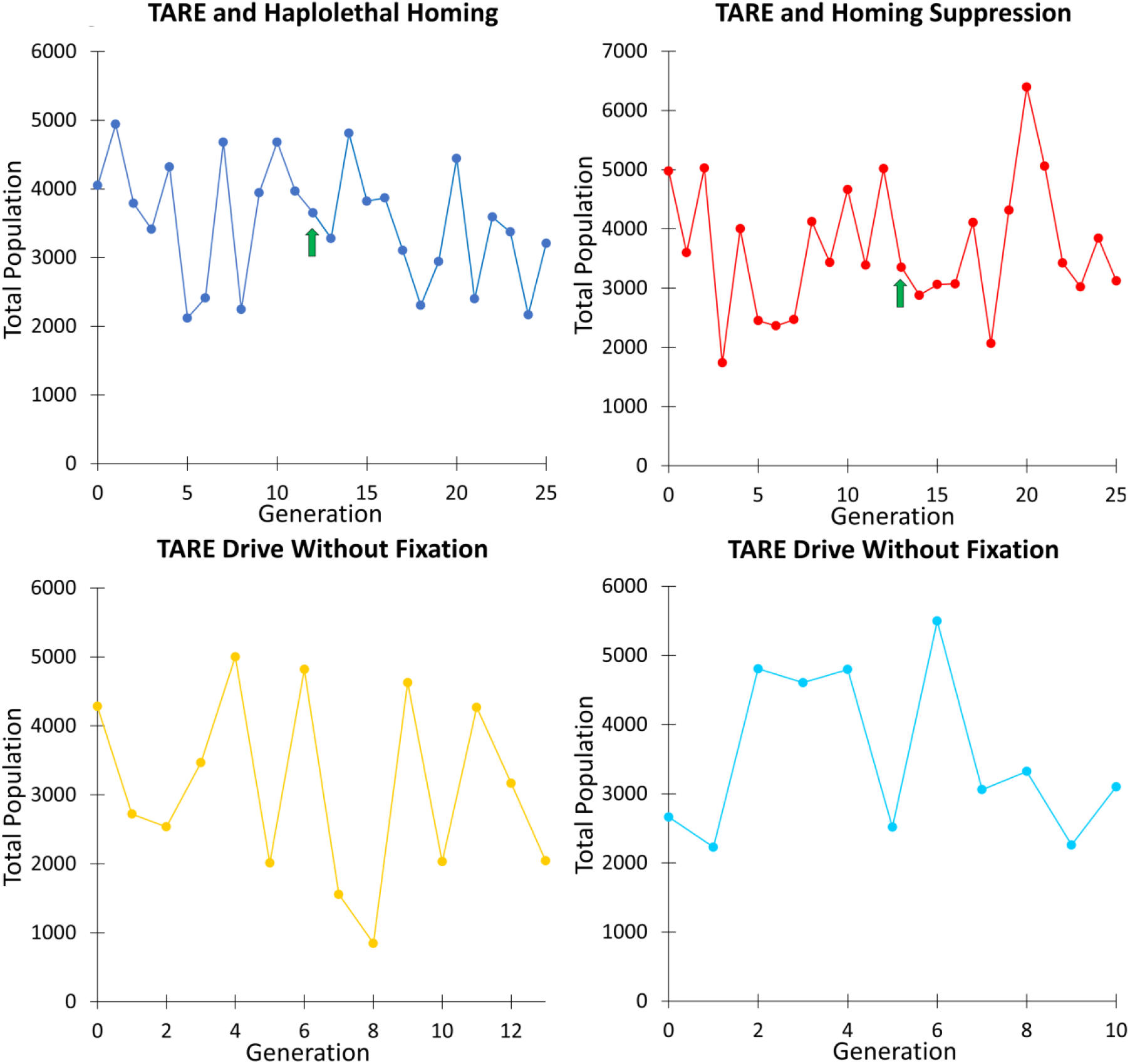
Population Sizes for Cage Study. The population is shown for each generation. Arrows indicate generations where homing drives were released in the first two cages.

**Table S1.** Comparison of different gene drive types.

| Drive type | High power suppression? | Threshold | Target gene | Preferred cas9 promoter | Engineering difficulty | Specific challenges |
| --- | --- | --- | --- | --- | --- | --- |
| **Homing drive** | yes | zero | essential | germline | low | requires high drive conversion |
| **Split homing drive** | no | self-limiting | essential | germline | low | requires high drive conversion |
| **Daisy drive** | yes* | eventually self-limiting | multiple essential | germline | moderate? | multiple targets |
| **Y-linked X-shredder** | yes | zero | X-chromosome | germline/embryo | high | Y chromosome engineering |
| **Killer-rescue** | no | self-limiting | variable | variable | moderate | variable mechanism |
| **TARE** | no | low | essential | any | low | none |
| **2-locus TARE** | no | moderate | two essential | any | low? | two alleles |
| **1-locus 2-drive TARE** | no | high | two essential | any | low? | two alleles |
| **TADE** | yes | low-moderate | haplolethal | germline | moderate? | haplolethal target |
| **TADE underdominance** | yes | moderate-high | haplolethal | germline/embryo | moderate? | haplolethal target |
| *Wolbachia* | no | moderate | N/A | N/A | low | complex organism |
| **Tethered drive system** | yes | varies** | as above | germline | low | requires high drive conversion |

N/A - not applicable. \*High power suppression until some drive elements are no longer supported by earlier elements. \*\*Tethered drives have variable confinement based on their cas9 containing element. “?” indicates that engineering difficulty is an estimate. Specific challenges can be highly variable based on the target species.

**Table S2.** Combined maximum likelihood parameter estimates from cage populations. Log-likelihood shows a relative probability (higher values indicate a better model fit) AICc: Akaike information criterion, corrected (low values indicate a better match of the model without overfitting) Effective population size refers to the percent of census size, which varies with each generation (the average was 3491) Fitness values are for drive homozygotes (with multiplicative fitness per allele). A value of 1 is equivalent to wild-type fitness, and “1” indicates that the parameter was fixed

| **Fitness cost model** | **Log-likelihood** | **AICc** | **Effective population size** | **Mating fitness** | **Fecundity fitness** | **Viability fitness** |
| --- | --- | --- | --- | --- | --- | --- |
| **None** | 96.5 | -191.0 | 3.02% | 1 | 1 | 1 |
| **All** | 100.4 | -191.9 | 3.68% | 0.943 | 0.846 | 0.962 |
| **All with mating = fecundity** | 100.4 | -194.3 | 3.58% | 0.867 | 0.867 | 1.00 |
| **Mating and fecundity** | 100.8 | -194.9 | 3.64% | 1.00 | 0.754 | 1 |
| **Mating and viability** | 100.3 | -194.0 | 3.56% | 0.741 | 1 | 1.00 |
| **Fecundity and viability** | 100.8 | -194.9 | 3.64% | 1 | 0.7539 | 1.00 |
| **Mating** | 100.3 | **<u>-196.3</u>** | 3.57% | 0.7322 | 1 | 1 |
| **Fecundity** | 100.8 | **<u>-197.2</u>** | 3.64% | 1 | 0.754 | 1 |
| **Viability** | 100.0 | **<u>-195.7</u>** | 3.52% | 1 | 1 | 0.853 |
| **Mating = fecundity** | 100.4 | **<u>-196.6</u>** | 3.59% | 0.867 | 0.867 | 1 |

